# Single-cell characterization of human periodontal ligament (PDL) cells reveals uniformly mesenchymal populations in matched maxilla and mandible pairs from four donors

**DOI:** 10.64898/2026.09.03.749128

**Authors:** Chloe Radermacher, Robert Dzhanaev, Christian Hasberg, Sabine Neuss-Stein, Andrea Gorgels, Julia Moellmann, Nikolaus Marx, Anne Babler, Christoph Kuppe, Rogerio Bastos Craveiro, Michael Wolf, Ali Modabber, Willi Jahnen-Dechent, Isabel Knaup

## Abstract

The periodontal ligament (PDL) is a mechanically active connective tissue whose fibroblasts and resident progenitor cells drive tooth support, remodeling, and regeneration. Although PDL cell populations are known to be heterogeneous, the composition of expanded primary PDL cultures and the extent to which anatomical origin shapes their transcriptome remain incompletely defined. Here we applied single-cell RNA sequencing to cultured primary PDL cells isolated from matched upper (maxilla) and lower (mandible) jaw sites of four donors, yielding 43,423 cells that were integrated across donors and analyzed by unsupervised clustering. The expanded cultures were uniformly mesenchymal and fibroblastic, essentially devoid of endothelial and immune signatures, and resolved into ten functionally distinct fibroblast substates rather than discrete cell types, including proliferating, collagen- and extracellular-matrix-high, contractile, and progenitor-like populations. Donor identity was the dominant source of transcriptional variation, with several substates restricted to individual donors. A donor-controlled comparison of maxilla versus mandible revealed only a small set of differentially expressed genes, headed by craniofacial positional-identity transcription factors (PITX1 higher in the mandible; BARX1, ALX1, and NKX6-1 higher in the maxilla), indicating that cultured PDL cells retain embryonic positional memory. In contrast, curated wound-healing gene modules did not differ significantly between jaws, and the transcriptional data therefore did not support the clinical impression of faster healing in the maxilla, suggesting that the healing asymmetry more likely reflects vascular, mechanical, and inflammatory signaling factors acting beyond the steady-state transcriptome. These findings characterize the cellular composition of expanded PDL cultures and identify positional identity as the principal jaw-dependent transcriptional feature.

## Introduction

The periodontal ligament (PDL) is a specialized fibrous connective tissue that anchors the tooth root to the alveolar bone, maintaining a remarkably consistent width despite widely varying mechanical loads, a feature that implies the existence of tightly regulated systems governing localized matrix resorption and synthesis ^1^. Within this tissue, the PDL fibroblast occupies a central functional position: cells are aligned parallel to collagen fibrils and establish direct membrane-to-fibril contacts through integrin-mediated linkages that transduce mechanical strain into intracellular signaling cascades ^1^. During active tissue formation, PDL fibroblasts display prominent rough endoplasmic reticulum, elaborate Golgi apparatus, and abundant secretory granules, reflecting a high capacity for matrix protein synthesis and turnover, while cytoskeletal adaptations allow these cells to withstand a continuously mechanically active environment ^1, 2^. Consistent with this synthetic role, transcriptomic profiling of native human PDL tissue has confirmed strong expression of collagens I, III, and VI, non-collagenous matrix proteins such as periostin and osteonectin, and proteoglycans including asporin, lumican, decorin, and osteomodulin, alongside integrin β1, underscoring the molecular basis of the fibroblast-matrix interface described structurally by earlier work ^3^. Notably, when PDL fibroblasts are removed from their native tissue context and expanded in culture, expression of key marker genes such as osteopontin, asporin, periostin, and osteonectin declines, suggesting that in vitro culture conditions favor a less differentiated, more immature cellular phenotype relative to intact tissue ^3^. Together, these findings establish the PDL fibroblast as both a structural anchor and a dynamic, mechanoresponsive regulator of periodontal tissue homeostasis.

The periodontium plays a pivotal role in the healing of periodontal wounds ^1^ suggesting that the periodontium might contain cells suitable for transplantation and therapeutic reconstruction of tissues destroyed by periodontal disease ^4^. This seminal study showed that beyond its differentiated fibroblast population, the PDL was indeed harboring a resident population of multipotent postnatal stem cells potentially suited for regenerative medicine and tissue engineering. Seo et al. demonstrated that cells isolated from human PDL tissue by single-colony selection expressed the mesenchymal markers STRO-1 and CD146/MUC18, distinguishing them from dental pulp stem cells and bone marrow stromal cells, and that these periodontal ligament stem cells (PDLSCs) could differentiate into cementoblast-like cells, adipocytes, and collagen-producing cells in vitro, as well as regenerate cementum/PDL-like structures upon transplantation into immunocompromised animals ^4^. This foundational identification of a stem cell compartment within an easily accessible tissue source has since been corroborated using standardized mesenchymal stem cell criteria, with isolated PDL cells consistently shown to be positive for CD73, CD90, and CD105 and negative for CD34 and CD45 ^5, 6, 7^. A small pilot study showed that clinically compatible, serum-free isolation protocols could yield viable PDLSCs for autologous regenerative applications ^5^. In parallel, primary PDL cells retaining the mesenchymal marker profile have been used to dissect mechanosensitive signaling, with toll-like receptor 4 identified as a key mediator of AKT and MAPK phosphorylation in response to compressive forces simulating orthodontic tooth movement, linking stem/progenitor cell biology directly to the mechanically active tissue environment described above ^6, 8^. It has been emphasized, however, that cells isolated from the PDL and expanded in vitro—commonly termed PDL-derived mesenchymal stem/stromal cells—are phenotypically diverse and preparation-dependent, and should be distinguished from the bona fide tissue stem cells whose stemness was demonstrated by rigorous in vivo lineage-tracing and transplantation assays ^9^.

Despite this apparently uniform surface marker profile, both clonal and bulk transcriptomic approaches have revealed that PDL progenitor populations are functionally and molecularly heterogeneous rather than homogeneous. Clonal analyses have segregated PDL progenitors into osteoblastic/cementoblastic clones and fibroblastic clones, with the former showing differential upregulation of Cadherin and Wnt pathway genes, including WNT2, WNT16, and WIF1, indicating that lineage commitment potential is encoded transcriptionally prior to overt differentiation ^10^. In a complementary murine cloning approach, a sparsely distributed subpopulation of Zbp1-expressing PDL cells was identified as possessing markedly higher osteogenic potential, with CRISPR/Cas9-based loss- and gain-of-function experiments confirming a direct functional role for Zbp1 despite the absence of any distinguishing morphological or standard progenitor marker differences between high- and low-osteogenic clones ^11^. Differentiation-directed transcriptomic comparisons have similarly shown that TGF-β1-driven ligament-fibroblastic differentiation and BMP-7-driven cementoblastic differentiation of PDLSCs are accompanied by distinct sets of extracellular matrix and membrane-associated differentially expressed genes, including divergent integrin and cadherin repertoires, reinforcing that the PDL progenitor pool gives rise to molecularly distinguishable descendant populations ^12, 13^. Functional heterogeneity extends to closely related periodontal cell types as well, as PDL cells exhibit modestly greater osteogenic and osteoclast-inductive capacity than anatomically adjacent alveolar bone-derived cells despite sharing fibroblastic morphology and comparable mesenchymal marker expression ^14^. Additional layers of heterogeneity are evident in the differential responsiveness of PDL fibroblasts to exogenous stimuli, as FGF1 promotes short-term proliferation and, with prolonged exposure, induces osteogenic marker expression while modulating the inflammatory response to mechanical compression ^15^, and as cannabinoid receptor CB1 signaling enhances osteo/dentinogenic differentiation via p38 MAPK and JNK pathways and rescues differentiation capacity suppressed by inflammatory cytokines TNF-α and IFN-γ ^16^. Collectively, these studies indicate that PDL progenitor and fibroblast populations comprise functionally distinct subsets whose behavior is shaped by both intrinsic transcriptional programming and extrinsic microenvironmental signals.

The advent of single-cell RNA sequencing (scRNA-seq) has substantially extended these clonal and bulk observations by enabling unbiased, genome-wide resolution of cellular heterogeneity within intact PDL populations. Comparative single-cell profiling of dental pulp stem cells and PDLSCs has resolved each population into three distinct clusters, with PDLSCs comprising predominantly osteogenic and myofibroblastic subpopulations, revealing cell-level heterogeneity not captured by bulk approaches ^17^. Similarly, scRNA-seq analysis identifying seven clusters across dental pulp and periodontal ligament stem cells showed that PDLSCs display a higher proportion of cells in G1 phase, indicative of lower proliferative capacity, together with trajectory analyses supporting specialized differentiation routes from a stem-like cluster toward more fibroblast-like states ^18^. Within intact murine PDL tissue, scRNA-seq has identified twelve cellular clusters comprising mesenchymal cells and macrophages, with the transcription factors Mkx and Scx partitioning mesenchymal cells into proteoglycan/elastic fiber-producing and collagen-producing subtypes, respectively, and with loss of Mkx driving pathological ossification and inflammation ^19^. In human periodontal tissue affected by periodontitis, single-cell transcriptomics has further distinguished IL-1β-expressing and RANKL-expressing PDL fibroblast subpopulations occupying distinct anatomical niches, both contributing to osteoclastogenic signaling through separate mechanisms ^20^. Extending this approach across periodontal tissue types, scRNA-seq comparing epithelium, gingiva, and PDL identified eight clusters overall, with a dominant PDL-specific cluster present across donors and developmental trajectory analyses suggesting that PDL cells represent a terminal differentiation stage relative to epithelial and gingival lineages ^21^. Together, these single-cell studies confirm that the PDL comprises multiple transcriptionally and functionally distinct subpopulations whose relative proportions and identities are relevant to both tissue homeostasis and disease. Single-cell profiling of in vitro-cultured PDL- and gingiva-derived MSCs from paired tissues of the same donors has likewise resolved tissue-specific and tissue-spanning subpopulations, while indicating that tissue-of-origin variability is comparatively limited and restricted to a subset of cellular processes ^22^. In intact murine PDL, a Plap-1 lineage-tracing atlas has resolved the native tissue into fibroblastic PDL cells alongside Ibsp+ cemento-/osteoblasts, Nes+ mural cells, S100B+ Schwann cells, and CD45+, CD31+, and Epcam+ non-stromal populations, and confirmed by direct lineage tracing that PDL cells give rise to osteoblasts and cementoblasts in vivo ^23^.

Beyond intrapopulation heterogeneity, accumulating evidence indicates that PDL cell biology also varies according to anatomical site, specifically between the maxilla and mandible. Comparative analysis of paired maxillary and mandibular PDLSCs from the same donors has shown that maxillary cells display greater proliferative capacity than their mandibular counterparts, along with higher osteogenic gene expression (ALP, RunX2, osteocalcin) and greater chondrogenic marker expression (SOX9, ACAN), despite both populations meeting standard mesenchymal stem cell marker criteria ^7^. No corresponding difference was observed in adipogenic differentiation potential, suggesting that jaw-of-origin differentially affects specific rather than global differentiation programs ^7^. At the signaling level, kinomic profiling of paired maxillary and mandibular PDLSCs from the same donors has revealed intrinsic differences in kinase activity and gene regulatory network engagement, with stronger EphA receptor expression in mandibular cells potentially inhibiting osteogenic differentiation, and greater PI3K-Akt pathway activation in mandibular relative to maxillary PDLSCs ^24^. These molecular distinctions help explain long-recognized clinical observations of faster wound healing and more favorable regenerative outcomes in the maxilla compared to the mandible, suggesting that jaw-specific intrinsic cellular programs, in addition to anatomical or vascular factors, may underlie these differences ^25^.

Taken together, the existing literature establishes that PDL cell populations are heterogeneous at both the clonal and single-cell level, and that this heterogeneity is further modulated by anatomical site of origin within the jaw. However, no study to date has applied single-cell transcriptomic profiling specifically to cultured PDL cells obtained from paired maxillary and mandibular sites within the same donors across multiple individuals, an approach that would allow simultaneous resolution of intrinsic subpopulation structure and jaw-associated transcriptional programs while controlling for donor-specific variability. Addressing this gap is essential for determining whether the site-dependent differences in proliferation, differentiation potential, and kinase signaling reported previously reflect shifts in the relative abundance of shared subpopulations, the presence of jaw-specific cell states, or both. The present study therefore aimed to perform single-cell RNA sequencing of cultured PDL cells derived from paired maxilla and mandible samples of multiple donors, to characterize the cellular heterogeneity of cultured human PDL populations and to determine whether maxillary and mandibular PDL cells harbor distinct, jaw-specific transcriptional programs.

## Materials and Methods

### Cell culture

Primary periodontal ligament cells (PDL), isolated from human third molars as previously described ^7^, were routinely maintained at 37°C in a humidified atmosphere with 5% CO₂. Cells were cultured in High Glucose DMEM (Gibco, Cat# 11965) supplemented with 10% FBS, 50mg/L L-ascorbic acid (Sigma-Aldrich), 100 U/mL penicillin, 100 µg/mL streptomycin. For routine passaging, cells were harvested at approximately 80% confluence using Trypsin-EDTA (Gibco, Cat# 25200056) and subcultured at a 1:5 ratio. The culture medium was refreshed every 48 hours. Cells were derived from matched maxilla (mx)- and mandible (mn) pairs from four donors Sp1, 3, 5, 12.

### Ethical statement and cell provenance

This study was conducted in accordance with the Declaration of Helsinki and was approved by the Ethics Committee of the Medical Faculty of RWTH Aachen University, approval no. EK374/19. Human PDL cells were isolated from third molars extracted for clinical indications from maxilla and mandibula in adult patients. All samples were fully anonymized prior to processing as Sp_x, and no personally identifiable information or clinical metadata linked to the donors was retained or used in this study. Written informed consent was obtained from all donors prior to tooth extraction.

### Single cell preparation and sequencing

Periodontal ligament cells were seeded from stocks of P1-2 cells (1-2 passage/expansion cycles) in six-well plates, grown to 80% confluency, and harvested by trypsinization, washed with phosphate-buffered saline containing 0.04% bovine serum albumin, and resuspended as a single-cell suspension. Cell concentration and viability were determined using a CASY® Cell Counter (Roche Innovatis AG, Mannheim, Germany), and only suspensions with a viability of >95% were used for library preparation. Cell suspensions were adjusted to the concentration recommended by the manufacturer, with a target recovery of approximately 1,600 cells per sample.

Single-cell 3′ gene-expression libraries were generated using the Chromium Next GEM Single Cell 3′ Kit v3.1 and the Chromium Controller (10x Genomics, Pleasanton, CA, USA) according to the manufacturer’s instructions. Individual cells were partitioned into Gel Beads-in-emulsion (GEMs), in which cells were lysed and polyadenylated RNA molecules were reverse-transcribed. During reverse transcription, complementary DNA molecules were labeled with cell-specific barcodes and unique molecular identifiers. Following GEM disruption, barcoded cDNA was recovered, purified, and amplified according to the protocol. Sequencing libraries were subsequently constructed by enzymatic fragmentation, end repair, A-tailing, adaptor ligation, and sample-index PCR.

Library concentration and fragment-size distribution were assessed using QuantiFluor® ONE dsDNA System (Promega Corporations, Fitchburg, WI USA) and a TapeStation HS D5000 (Agilent Technologies, Santa Clara, CA, USA). Libraries were pooled and sequenced on a NovaSeq 6000 (Illumina, San Diego, CA, USA) using paired-end sequencing according to the read configuration recommended for the Chromium Single Cell 3′ v3.1 chemistry. A minimum sequencing depth of approximately 20,000 read pairs per cell was targeted.

Raw base-call files were demultiplexed, and sequencing reads were aligned to the human reference genome (GRCh38, reference version) using Cell Ranger (10x Genomics). Gene–barcode matrices were generated using the Cell Ranger count pipeline and used as input for subsequent quality control and single-cell transcriptomic analyses.

### Curing of single-cell RNA sequencing data

The computational analysis of single-cell RNA sequencing (scRNA-seq) data was performed using the Scanpy framework ^26^ (v1.9.3) implemented in Python (release 3.11). Computations were performed using the high-performance computing cluster HPC of RWTH Aachen University under project *rwth1717*.

Initial input consisted of filtered gene-barcode matrices generated by the Cell Ranger pipeline (10x Genomics), corresponding to high-quality transcriptomic profiles of single cells. These matrices include barcodes that passed the Cell Ranger quality control metrics and therefore represent putative viable cells. Following data loading, an initial filtering step was carried out to remove cells with fewer than 200 expressed genes, to eliminate barcodes likely associated with low-quality or empty droplets.

To eliminate low-quality data, cells exhibiting more than 20% of their total reads mapping to mitochondrial genes were excluded. Furthermore, an upper threshold was applied to filter out potential multiplets, which are artifacts arising from the co-encapsulation of more than one cell within a single droplet. Specifically, this upper bound was defined as the 98^th^ percentile of the distribution of the number of genes detected per cell (i.e., genes with nonzero counts). Cells exceeding this threshold were removed to reduce the impact of outlier transcriptomes that may represent multiplets distorting downstream analyses.

To further address the issue of doublets, the scvi-tools ^27^ (v1.3.3) package was employed. Specifically, the SCVI model was used as a probabilistic framework to model transcriptomic variability and identify likely doublet populations in a data-driven manner. This enabled removal of doublets by leveraging a latent representation of the transcriptome learned through variational inference, without relying on predefined marker genes or cell type labels.

Following quality control and doublet exclusion, the data were concatenated across all samples and subsequently subjected to normalization and transformation. Total-count normalization was performed to account for differences in sequencing depth across cells, by scaling each cell’s transcript count to a target sum of 10,000 counts. Normalized data were then logarithmically transformed, and the untransformed data were stored as a .raw attribute for potential later use. These preprocessing steps ensure comparability of gene expression levels across cells and stabilize variance for downstream statistical analyses.

### Single-cell RNA-seq analysis pipeline

Data were analyzed using the annotated data set computed as described above (integrated_matrix_PDL_CR_with_distances.h5ad) as input for a generic Anthropic Claude Science® single cell sequencing pipeline. The pipeline was running Python scripts on an Apple MacStudio Computer equipped with an Apple M2 Ultra processor and 64 GB RAM. The data set comprised expression data from 43,423 cells × 28,982 genes representing four donors: Sp1 (12,510), Sp3 (13,302), Sp5 (6,484), Sp12 (11,127) with maxilla/mx (24,291) and mandible/mn (19,132), respectively resulting in 8 samples (jaw × donor) altogether.

The input .h5ad file arrived pre-processed as described above: QC, doublet removal (0 remain), total-count normalization + log1p (in X), raw counts (layers[“counts”]), scVI integration (X_scVI, 10 dims), a UMAP embedding, and a k-NN graph (n_neighbors=15, on scVI space). This pipeline ran the downstream stages only: clustering, composition, marker detection, annotation — no re-QC, re-normalization, or re-integration.

## Results

### Clustering

Leiden clustering of the existing scVI neighbor graph of the input annotated data file was replotted at three resolutions: 0.2 → 5 clusters, 0.5 → 10 clusters (primary), 1.0 → 17 clusters. Resolution 0.5 resolves the biologically distinct substates without over-fragmenting the dominant fibroblast body. Figure 1 shows the three UMAP embeddings side by side; the 10-cluster resolution 0.5 solution is used for all downstream analyses.

**Figure 1.**
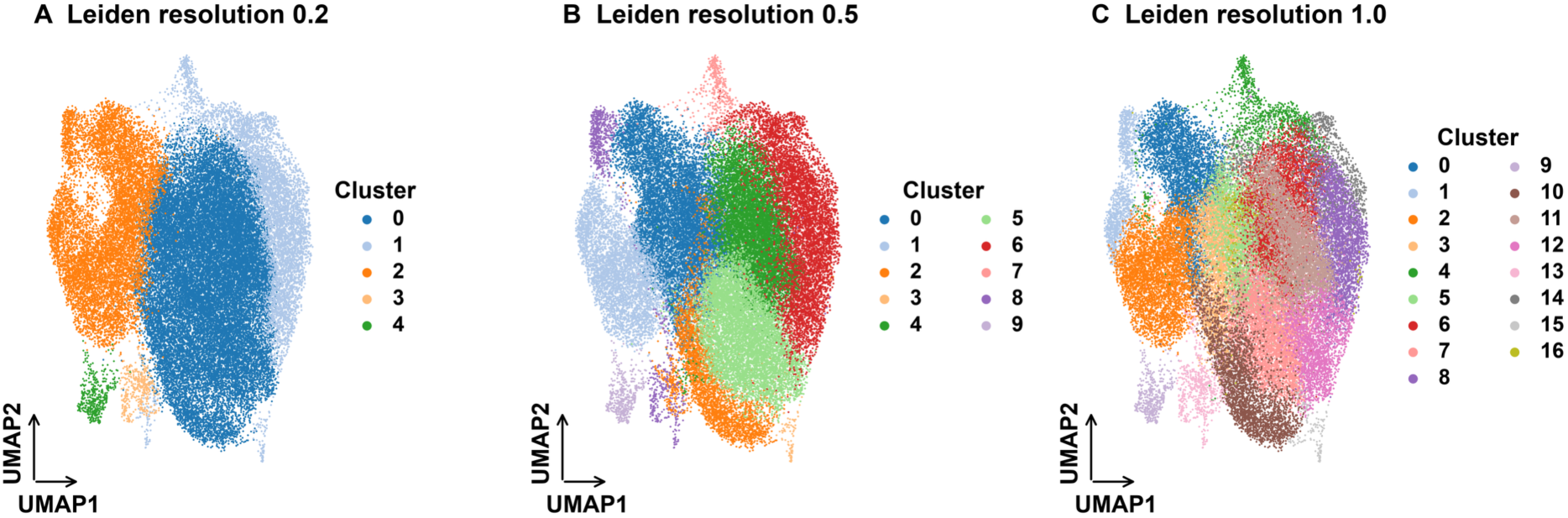
UMAP at three Leiden resolutions. Three UMAP embeddings of all 43,423 PDL cells, each point representing one cell, colored by Leiden cluster at (A) resolution 0.2 (5 clusters), (B) resolution 0.5 (10 clusters) and (C) resolution 1.0 (17 clusters). Cluster colors are arbitrary identity labels (see legend at right of each panel); increasing resolution splits the data into progressively finer clusters. Resolution 0.5 (clusters C0–C9) was selected for all downstream analyses. UMAP, Uniform Manifold Approximation and Projection — a non-linear 2-D projection of the scVI latent space; axes are arbitrary and carry no units. Cluster color mapping — C0 blue, C1 light-blue, C2 orange, C3 light-orange, C4 green, C5 light-green, C6 red, C7 pink, C8 purple, C9 light-purple.

### UMAP overview

Clusters are spatially coherent and donor-mixed across the main body (scVI integration is effective). QC metrics (library size, gene count, mito%, ribo%) are largely uniform across clusters — no cluster is a QC artifact. Figure 2 shows the eight-panel UMAP overview, in which the clusters are spatially coherent and donor-mixed, and the quality-control metrics vary little between clusters.

**Figure 2.**
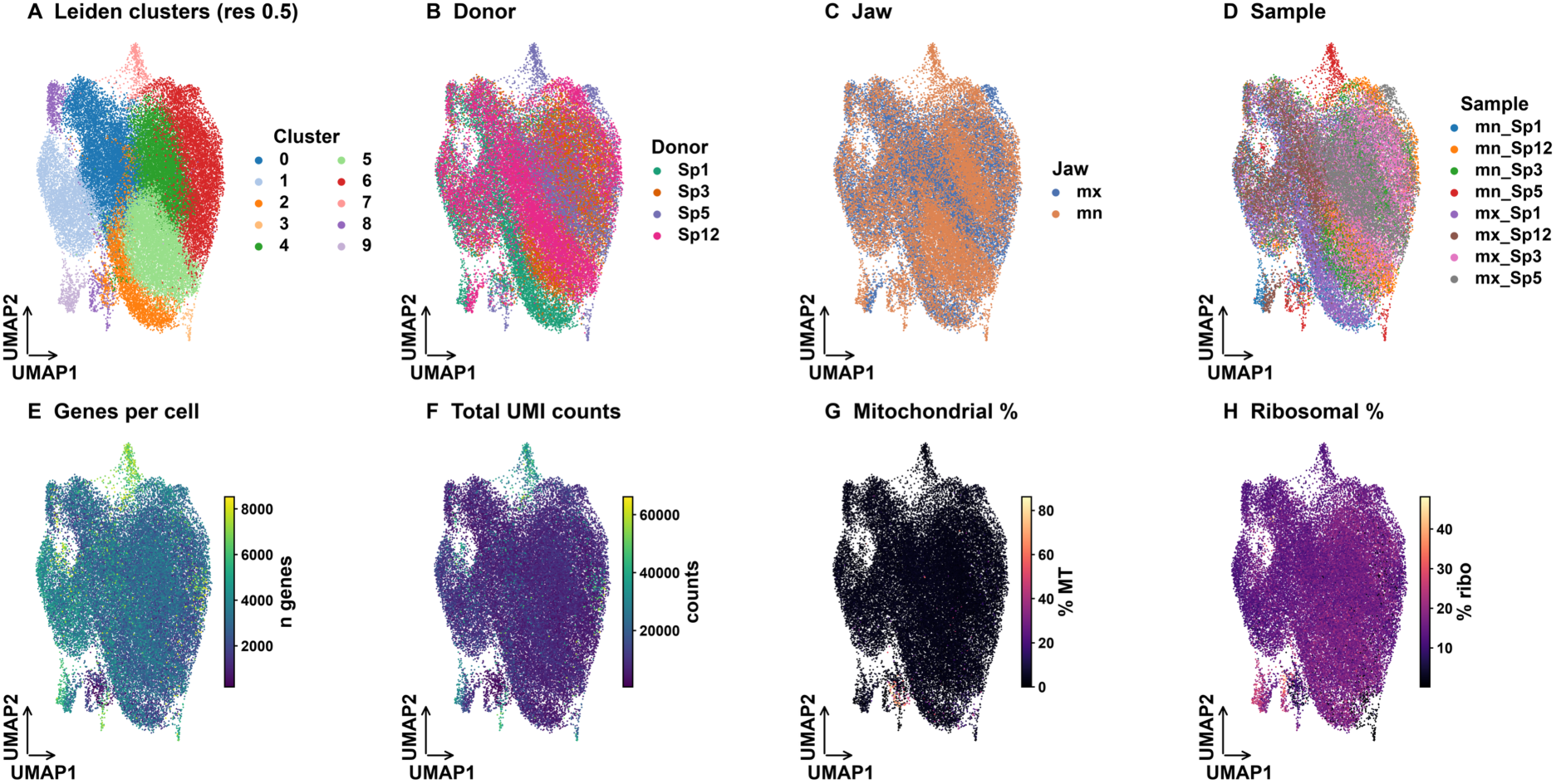
UMAP overview: clustering, sample identity and quality-control metrics. Eight UMAP panels of the same embedding. Top row (categorical, one color per group): (A) Leiden clusters C0–C9, (B) Donor (Sp1, Sp3, Sp5, Sp12), (C) Jaw, and (D) Sample (the eight donor×jaw combinations). Bottom row (continuous metrics, colour bar at right of each panel): (E) genes detected per cell and (F) total UMI counts (viridis scale), (G) mitochondrial content (% MT) and (H) ribosomal content (% ribo) (magma scale). Each point is one cell. Abbreviations: mx, maxilla; mn, mandible; QC, quality control; UMI, unique molecular identifier; MT, mitochondrial; ribo, ribosomal.

### Cell-type annotation

Marker-set scoring shows a uniformly mesenchymal population — endothelial (PECAM1/VWF/CDH5) and immune (PTPRC/CD68/CD3E) signatures score ≈ 0 in every cluster. This is the expected profile of an expanded primary PDL cell culture in base medium. The 10 clusters therefore represent functional PDL/fibroblast substates, not distinct cell lineages. Two proliferating clusters were confirmed by cell-cycle scoring (cluster 1: 99% S/G2M; cluster 8: 69% G2M/M). Figure 3 shows the annotated substates and their cell-cycle phase, and Figure 4 shows each cluster’s size alongside its fraction of cycling (S+G2M) cells, confirming clusters 1 and 8 as the proliferative substates.

**Table 1.** Cell-type annotation.

| Cluster | Cell-type label | N cells | % total | Dom. donor | Dom. jaw | % cycling |
| --- | --- | --- | --- | --- | --- | --- |
| 0 | Stress/MT-high fibroblast | 10437 | 24.0 | Sp1 (42%) | mx (68%) | 42.0 |
| 1 | Proliferating fibroblast (S/G2M) | 4420 | 10.2 | Sp1 (47%) | mx (60%) | 99.0 |
| 2 | Contractile fibroblast (Sp1) | 3558 | 8.2 | Sp1 (86%) | mx (60%) | 8.0 |
| 3 | ECM-high fibroblast (Sp5/mn) | 84 | 0.2 | Sp5 (86%) | mn (90%) | 5.0 |
| 4 | Collagen-high fibroblast | 9018 | 20.8 | Sp3 (58%) | mx (56%) | 4.0 |
| 5 | ECM/decorin fibroblast | 7502 | 17.3 | Sp12 (43%) | mn (67%) | 4.0 |
| 6 | VCAN+ progenitor-like fibroblast | 6682 | 15.4 | Sp3 (50%) | mx (63%) | 3.0 |
| 7 | Myofibroblast/elastic (Sp5/mn) | 413 | 1.0 | Sp5 (96%) | mn (99%) | 9.0 |
| 8 | Proliferating fibroblast (G2M/M) | 869 | 2.0 | Sp1 (34%) | mn (51%) | 69.0 |
| 9 | Ribosome-high fibroblast | 440 | 1.0 | Sp12 (52%) | mx (53%) | 50.0 |

**Figure 3.**
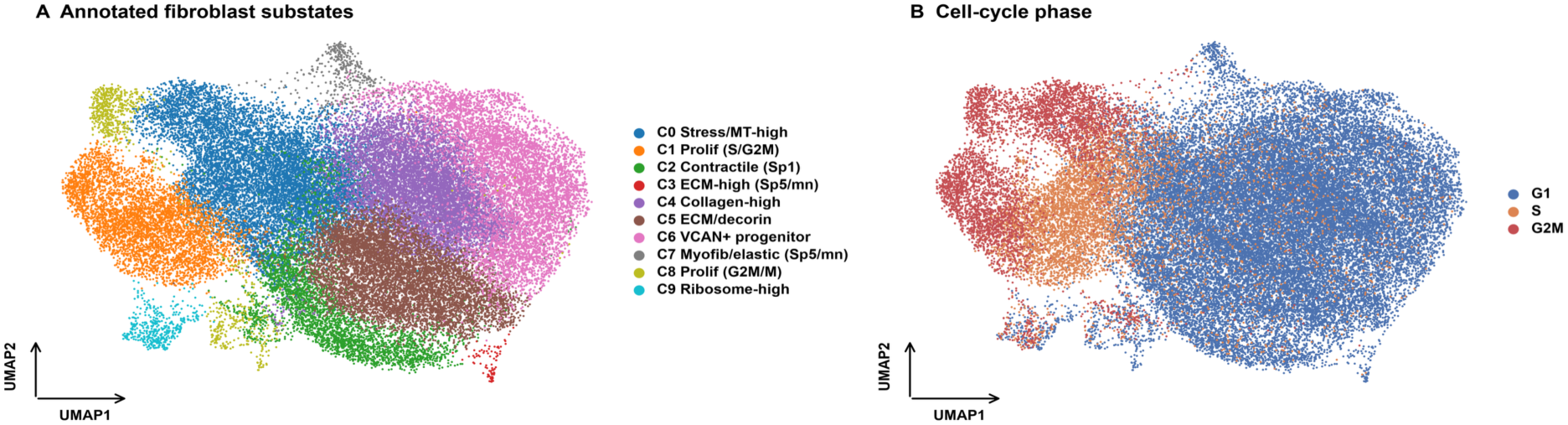
Annotated UMAP and cell-cycle phase. (A) UMAP colored by annotated fibroblast substate (clusters C0–C9); the legend gives the short functional label of each substate (e.g. C0 Stress/MT-high, C1 Proliferating (S/G2M), C5 ECM/decorin, C7 Myofibroblast/elastic). (B) The same UMAP colored by inferred cell-cycle phase: G1 (blue), S (orange), G2M (red). Each point is one cell. All fibroblast substates share a common mesenchymal identity — the labels describe functional states, not distinct cell types. Abbreviations: mx, maxilla; mn, mandible; MT, mitochondrial.

**Figure 4.**
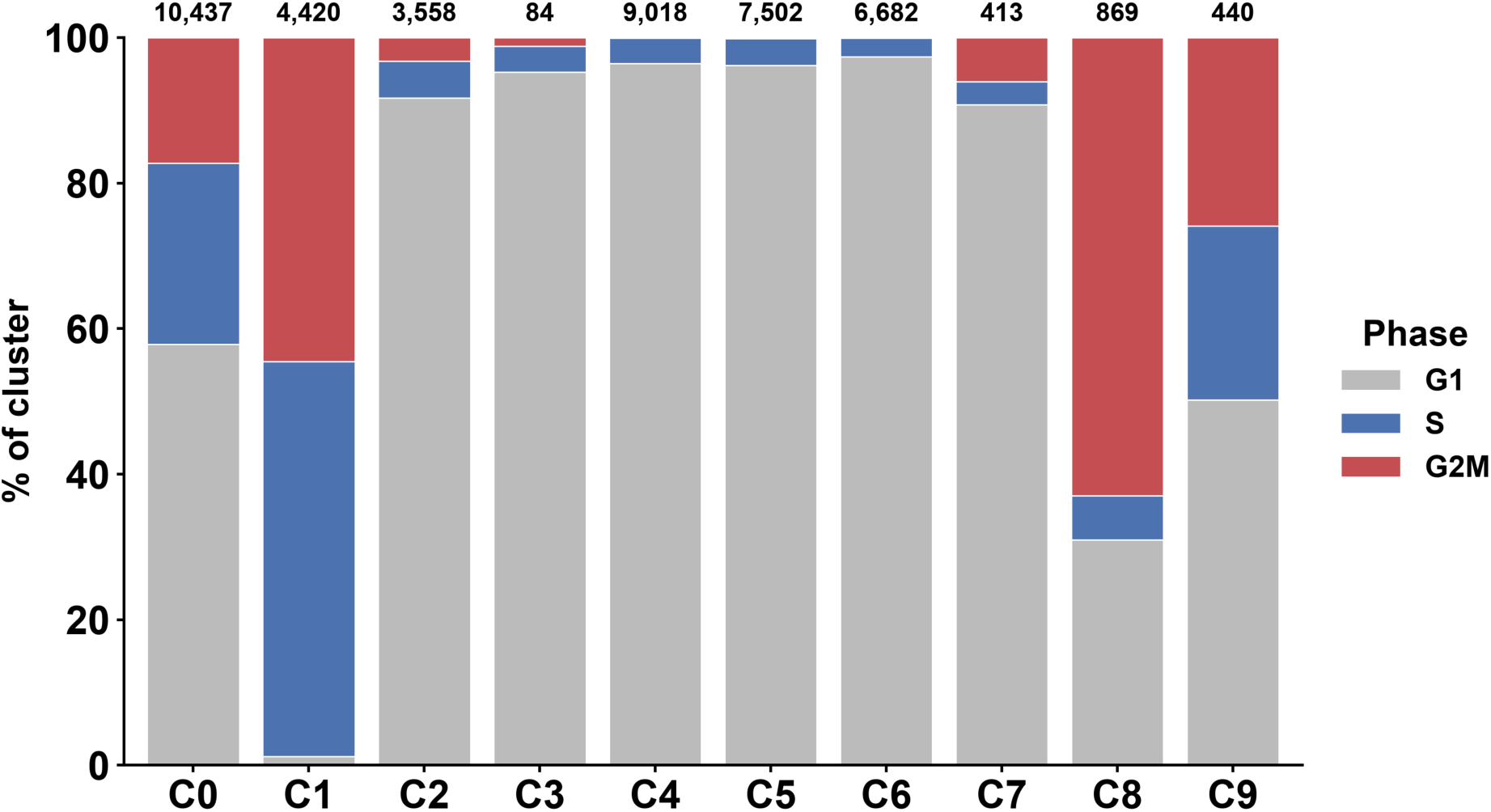
Cell-cycle phase composition and proliferative activity per cluster. Each bar is one cluster (C0–C9); segments give the percentage of that cluster’s cells in each inferred cell-cycle phase — G1 (grey), S (blue), G2M (red) — stacked to 100 %. The Y-axis gives the percentage of the cluster’s cells in each phase; the cluster size (n cells) is printed above each bar. Clusters C1 (99 % cycling) and C8 (69 %) are the proliferating populations, whereas C3–C6 are largely quiescent (G1). Leiden cluster numbers reflect the community-detection ordering of the shared-neighbor graph, not cluster size: for example, C3 (ECM-high, Sp5/mn; n = 84) is far smaller than C4 (collagen-high; n = 9,018) despite its lower index, because C3 is a small donor- and jaw-restricted community that the algorithm resolved as its own cluster.

### Cluster composition

Most clusters draw from all four donors and both jaws. Notable exceptions: cluster 2 (contractile, MYLK/PDE5A) is 86% Sp1; clusters 3 and 7 (ECM-high / elastic-myofibroblast) are Sp5-specific and mandible (mn)-specific (86–96% Sp5, 90–99% mn) — small populations (84 and 413 cells) that may reflect a donor-specific state or residual technical effect in the smallest sample (mn_Sp5, 1,153 cells). Figure 5 shows the donor, jaw and sample composition of every cluster as stacked bars, while Figure 6 shows cluster abundance resolved by jaw (A) and by donor (B), highlighting the Sp1-dominated cluster 2 and the Sp5/mn-restricted clusters 3 and 7.

**Figure 5.**
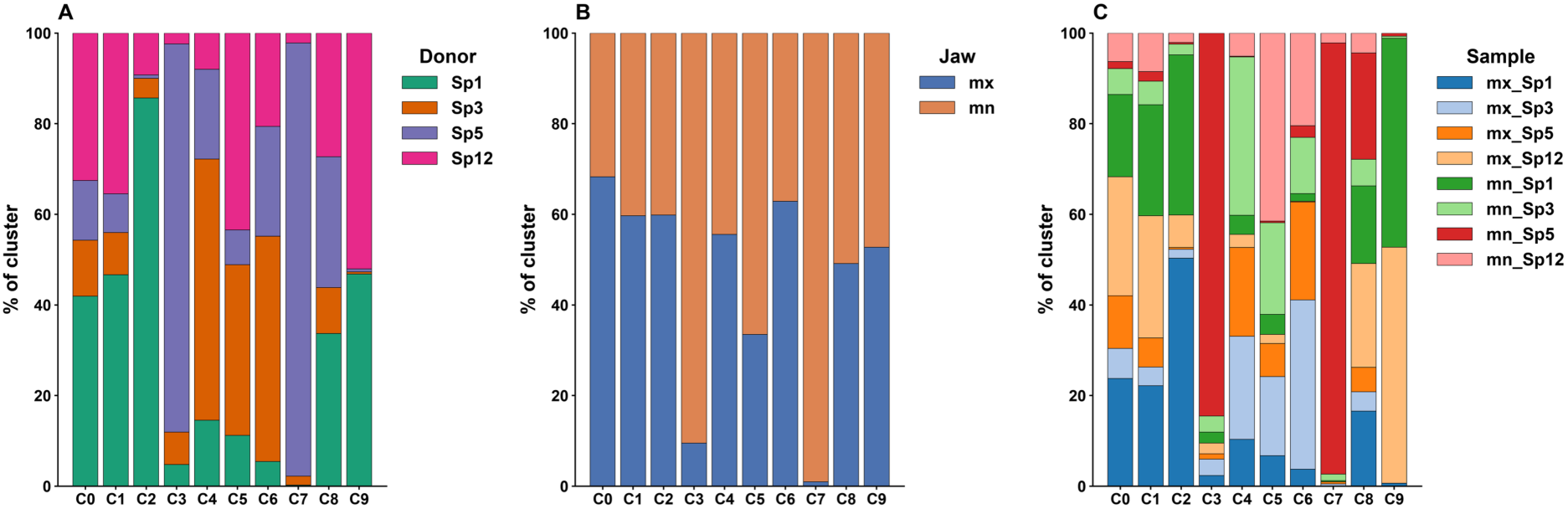
Cluster composition by donor, jaw and sample. Three 100 %-stacked-bar panels; each bar is one cluster (C0–C9) and segment heights give the percentage of that cluster’s cells contributed by each group. (A) Donor: Sp1 (green), Sp3 (orange), Sp5 (purple), Sp12 (magenta). (B) Jaw: mx (blue, maxilla), mn (orange, mandible). (C) Sample: the eight donor×jaw combinations (legend at right).

**Figure 6.**
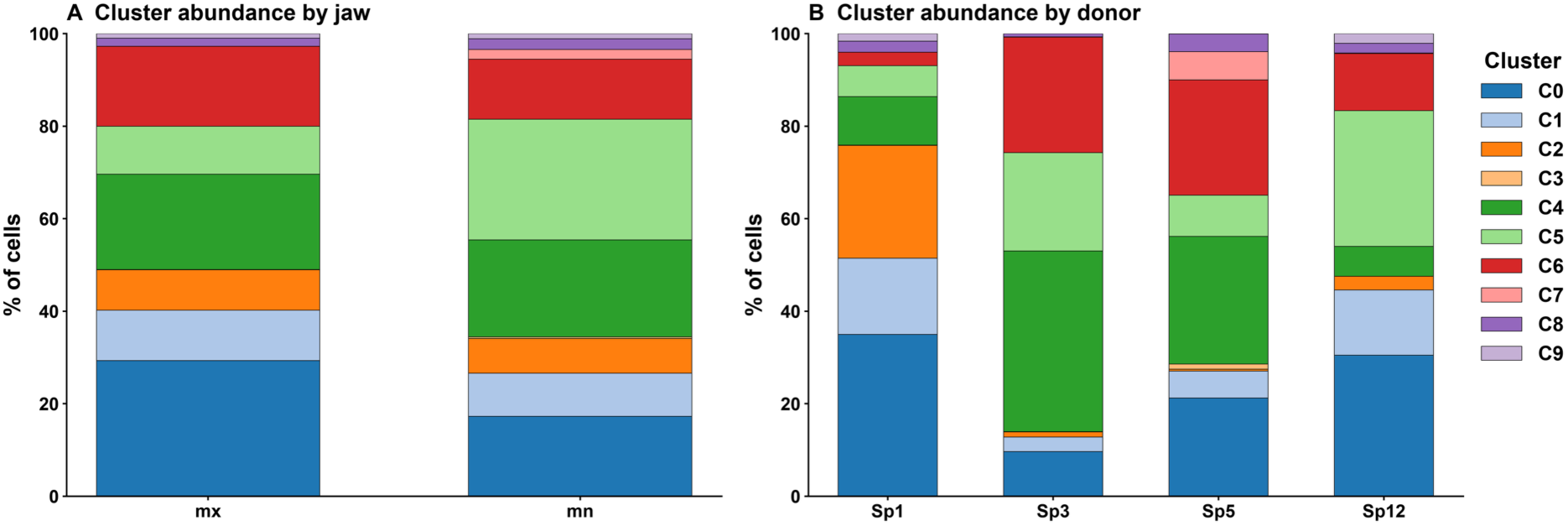
Cluster abundance by jaw and by donor. (A) Cluster composition within each jaw: each bar is one jaw (mx, maxilla; mn, mandible) and segments give the percentage of that jaw’s cells assigned to each cluster (C0–C9). (B) Cluster composition within each donor (Sp1, Sp3, Sp5, Sp12), same encoding. Colors follow the common cluster palette (legend at right). Cluster C0 (stress/MT-high) is enriched in the maxilla and C5 (ECM/decorin) in the mandible, but the dominant structure is donor-restricted — C2 (contractile) is almost exclusively Sp1, C5 is enriched in Sp12, and C7 (myofibroblast/elastic) in Sp5 — showing that donor identity, rather than jaw of origin, dominates cluster membership. mx, maxilla; mn, mandible.

### Marker genes

The full Wilcoxon rank-sum results (289,820 rows) are listed in supplementary STable 1 (markers_per_cluster.csv). Figure 7 shows the top-marker dot plot — the five most differentially expressed genes per cluster (Wilcoxon rank-sum test, each cluster versus all others) — with clean cluster-specific expression blocks. To place the clusters relative to published periodontal-ligament cell isolates, we scored each cell for canonical mesenchymal-stromal-cell (MSC) and PDL-associated gene programs (Figure 8).

**Figure 7.**
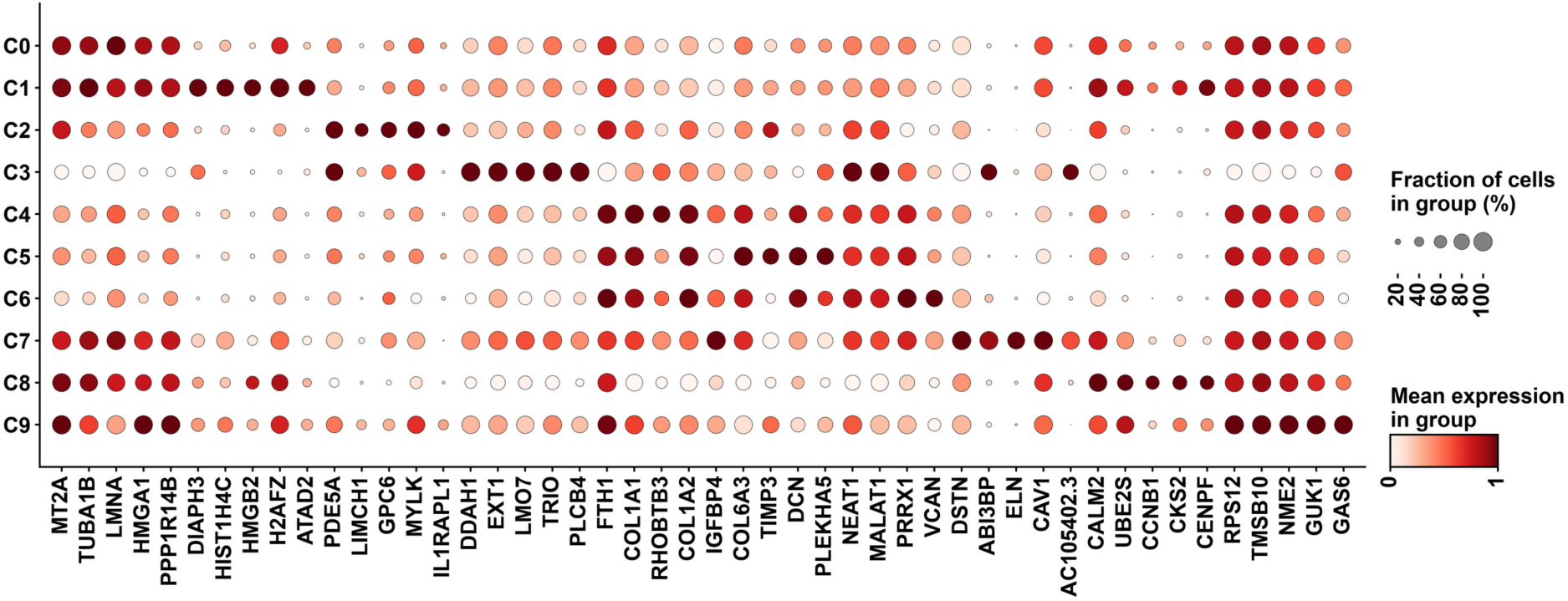
Top-marker dot plot. The five most differentially expressed genes per cluster (Wilcoxon rank-sum test, each cluster versus all others), grouped along the x-axis under the cluster they mark; rows are clusters 0–9. Dot color encodes mean expression scaled 0–1 per gene (Reds colormap); dot size encodes the fraction of cells in the cluster expressing the gene (size key at right).

All clusters share a uniformly mesenchymal identity — the ISCT MSC-negative (hematopoietic/endothelial) panel scores ≈ 0 throughout — and cluster C7 (myofibroblast/elastic, Sp5/mn) ranks highest on a composite MSC/PDL identity index, expressing the strongest ISCT surface markers (THY1/CD90, ENG/CD105) and the highest periostin (POSTN), followed by C9 (ribosome-high). These clusters most closely match the marker profile reported for cultured PDL-derived MSCs. Because all cells were culture-expanded in monolayer, this ranking reflects intrinsic in-vitro marker and program expression rather than a reference-based cell-type classification: C7 is the cluster whose cells intrinsically express the most typical MSC/PDL markers under these culture conditions, not a formally identified PDL-MSC population. Figure 8B and SFigure 1 show that cultured PDL cells retain low but heterogeneous expression of PLAP-1 / Asporin, a the PDL marker concentrated in C5, an ECM/decorin-high fibroblast state, consistent with a matrix subpopulation that may regulate mineralization as proposed by Yamada et al. ^28^. Retention of the ligament/PDL matrix marker (PLAP-1) together with near-absence of the cementoblast marker (CEMP1) and likely also cementum attachment protein (CAP) shown in SFigure 1 is consistent with the culture-expanded cells corresponding to the ligament-fibroblastic rather than the cementoblastic progenitor state described by Mun et al. ^12^.

**Figure 8.**
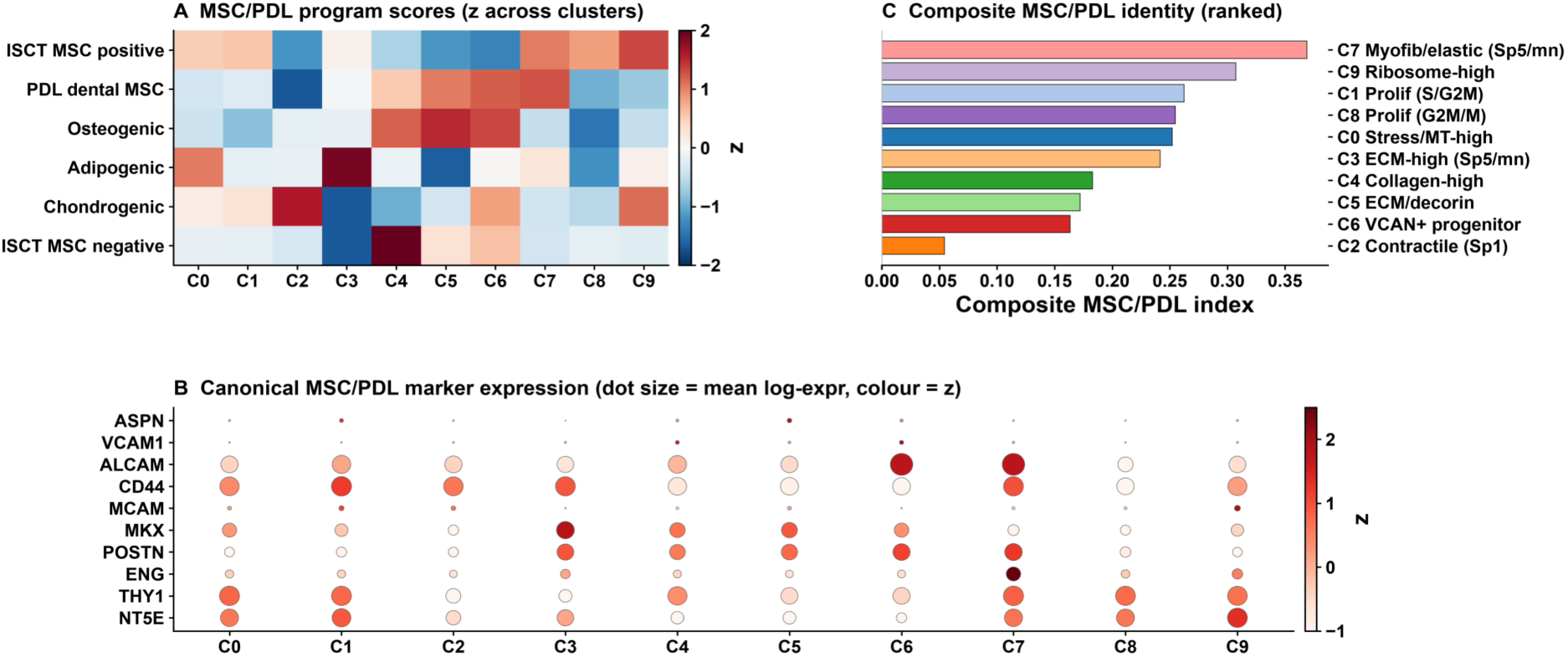
Mesenchymal (MSC) and periodontal-ligament (PDL) cell-program analysis per cluster. (A) Heatmap of mean per-cell program scores (sc.tl.score_genes on log-normalized expression) for six literature-defined gene sets — ISCT minimal MSC-positive markers (NT5E/CD73, THY1/CD90, ENG/CD105); the ISCT MSC-negative panel (PTPRC/CD45, CD34, CD14, CD19, HLA-DRA); a dental/PDL-MSC set (POSTN, MKX, SCX, MCAM/CD146, CD44, ALCAM/CD166, NES, VCAM1, ASPN, TNMD, S100A4); and osteogenic, adipogenic and chondrogenic differentiation programs — z-scored across clusters (red, above the cross-cluster mean; blue, below). (B) Dot plot of the canonical MSC/PDL surface and matrix markers (dot size = mean log-expression, color = z-score across clusters). (C) Composite MSC/PDL identity index (mean of the ISCT-positive and PDL-MSC program scores minus the MSC-negative score), ranked by cluster; bars are color-matched to the cluster palette. The ISCT MSC-negative panel scores ≈ 0 in every cluster, confirming a uniformly mesenchymal population free of hematopoietic/endothelial contamination. Cluster C7 (myofibroblast/elastic, Sp5/mn) ranks highest on the composite index — carrying the strongest ISCT surface markers (THY1/CD90, ENG/CD105) and the highest periostin (POSTN) — followed by C9 (ribosome-high); these clusters most closely match the canonical PDL-derived MSC profile reported for cultured PDL isolates. This composite ranking reflects intrinsic in-vitro marker/program expression among culture-expanded cells, not a reference-based cell-type call.

**SFigure 1.**
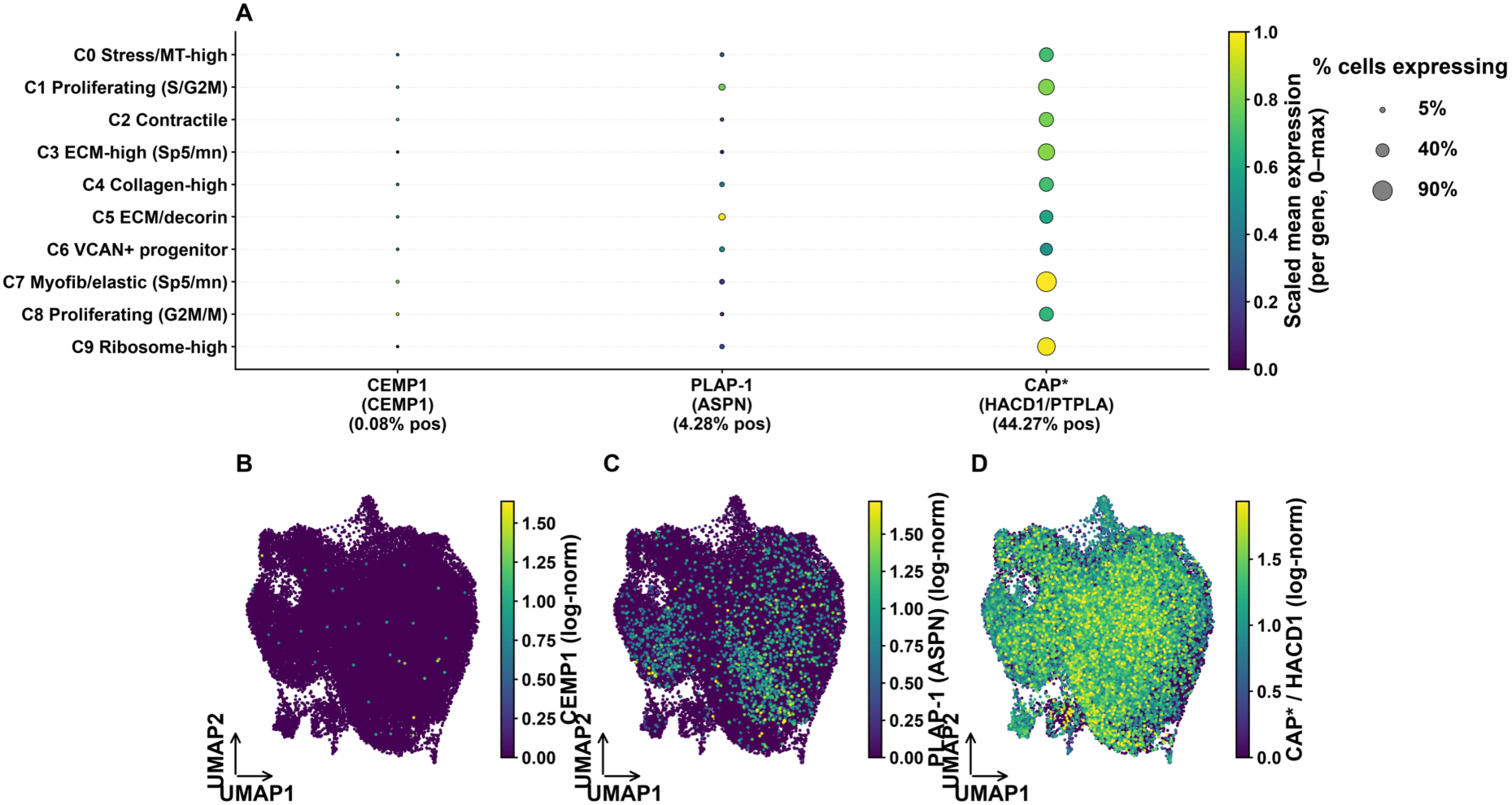
Expression of cementum and PDL matrix markers (CEMP1, PLAP-1, CAP) in culture-expanded PDL cells. (A) Dot plot of CEMP1 (gene CEMP1), PLAP-1 (gene ASPN) and CAP (gene HACD1/PTPLA) across the ten Leiden cell states; dot size encodes the percentage of expressing cells and colour the per-gene mean expression scaled 0–1 (absolute detection is given in parentheses under each column label). (B–D) UMAPs of all 43,423 cells coloured by CEMP1, ASPN and HACD1 expression (log-normalized). The cementoblast marker CEMP1 is essentially undetectable (0.08 % of cells, 35 transcripts total, ≤1 per cell) with no state significantly enriched, whereas PLAP-1/asporin (ASPN) is detected in 4.3 % of cells overall (dataset mean 0.036 log-normalized, ≈ 0.05 raw counts per cell) — low but non-uniform expression concentrated in the ECM/decorin matrix state, with C5 (ECM/decorin) and C1 (proliferating, S/G2M) the only substates expressing ASPN significantly above the dataset mean (Mann-Whitney U with Benjamini-Hochberg correction). *Cementum attachment protein (CAP) is not an independent gene but a splice variant of HACD1/PTPLA (3-hydroxyacyl-CoA dehydratase 1/protein tyrosine phosphatase-like) that encodes a truncated 140-amino-acid PTPLa/CAP polypeptide (GenBank AY455942.1 / AAR22554.1) expressed in cementoblasts and cementoblast precursors ^13^. The scRNA-seq reference annotates HACD1 as a single gene-level feature and does not resolve this variant; the CAP lane therefore reports total HACD1/PTPLA (detected in 44.3 % of cells across all states, reflecting its housekeeping role in fatty-acid elongation) and cannot be equated with the CAP splice variant. Until this discrepancy is resolved at the transcript level, retention of the ligament/PDL matrix marker (PLAP-1) together with near-absence of the cementoblast marker (CEMP1) is consistent with the culture-expanded cells corresponding to the ligament-fibroblastic rather than the cementoblastic progenitor state described by Mun et al. ^12^.

SFigure 2 shows that expression of PDL-associated transcription factors MKX/mohawk was retained and graded along the matrix axis — highest in ECM-high, ECM/decorin, collagen-high; lowest in proliferating cells mirroring the Mkx-associated matrix program described in native PDL ^19^. In contrast SCX/scleraxis was largely absent with only a weak signal in the ribosome-high state, and ZBP1, which marks high-osteogenic murine PDL clones ^11^ was below the detection limit in all culture expanded PDL cells.

**SFigure 2.**
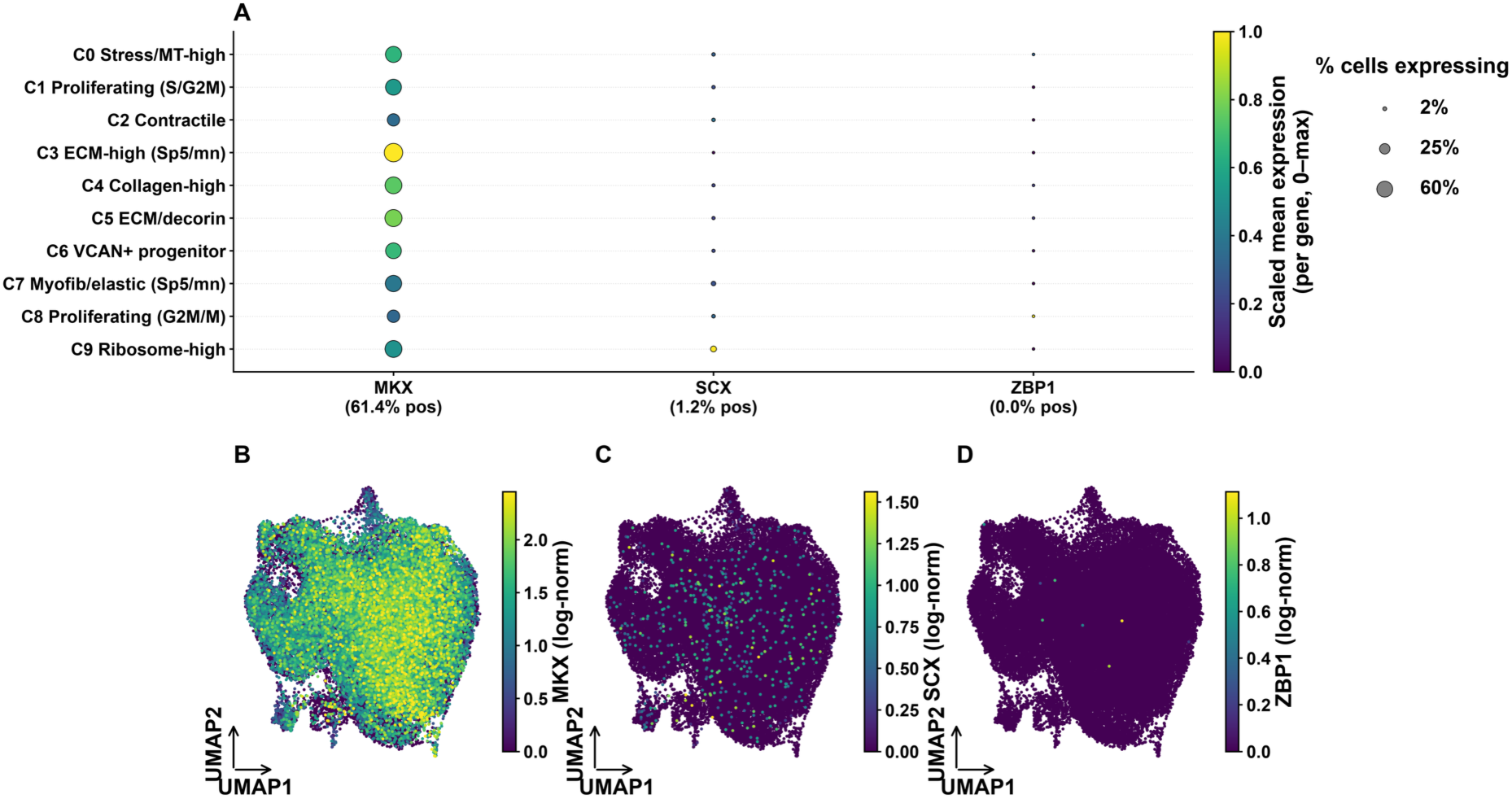
Expression of PDL/tendon-associated transcription factors (MKX, SCX, ZBP1) in culture-expanded PDL cells. (A) Dot plot of MKX (mohawk), SCX (scleraxis) and ZBP1 across the ten Leiden cell states. Dot size encodes the percentage of cells expressing the gene; colour encodes mean expression scaled 0–1 within each gene (per-gene absolute detection is given in parentheses under each column label). (B–D) UMAPs of all 43,423 cells coloured by MKX, SCX and ZBP1 expression (log-normalized). MKX is broadly retained (61.4 % of cells) and graded along the extracellular-matrix axis, being highest in the ECM-high, ECM/decorin and collagen-high fibroblast states and lowest in the proliferating fractions. In contrast, SCX is largely absent (1.2 % of cells, ≤ 2 transcripts per cell) and ZBP1 is essentially undetectable (8 transcripts across 43,423 cells). Statistical enrichment per state was assessed by Mann-Whitney U with Benjamini-Hochberg correction; values are tabulated in the supplementary per-state table.

Note that annotation labels are descriptive substates from marker enrichment + cell-cycle scoring, not reference-based calls. No PDL-specific CellTypist model was available. The Sp5/mn-restricted clusters (3, 7) are small and should be treated as donor-specific observations pending confirmation with additional Sp5 mandible material.

### Wound healing: maxilla vs mandible

The periodontium plays a pivotal role in the healing of periodontal wounds ^1^. Anecdotal clinical observation suggests that surgical wounds heal faster in the maxilla than the mandible. We asked whether the transcriptional programs associated with wound healing are elevated in maxillary PDL cells relative to mandibular cells in this dataset. The following section evaluates whether wound-healing-associated transcriptional programs differ between cells from the two jaws, using the same donor-paired statistical framework. These wound-healing-associated programs were defined as seven curated, fibroblast-relevant gene modules, each capturing one arm of the canonical wound-healing cascade and each corresponding to a specific KEGG ^29, 30^ and WikiPathways ^31^ pathway: (i) proliferation (signature genes MKI67, PCNA, TOP2A, CCNB1, CDK1, BIRC5; KEGG hsa04110 Cell cycle, WikiPathways WP179); (ii) angiogenesis (VEGFA, VEGFC, KDR, FLT1, ANGPT1/ANGPT2, PECAM1; KEGG hsa04370 VEGF signaling, WikiPathways WP1539 Angiogenesis); (iii) extracellular-matrix (ECM) remodeling by matrix metalloproteinases and their inhibitors (MMP1, MMP2, MMP9, MMP14, TIMP1–TIMP3, PLAU/PLAUR; KEGG hsa04512 ECM–receptor interaction, WikiPathways WP129 Matrix metalloproteinases); (iv) collagen and ECM synthesis (COL1A1, COL1A2, COL3A1, FN1, LOX, POSTN, ELN; KEGG hsa04974 Protein digestion and absorption); (v) growth-factor signaling and cell migration (TGFB1, TGFB3, PDGFA/PDGFB, FGF2, FGF7, HGF, CCN1/CCN2, HBEGF; KEGG hsa04350 TGF-β signaling, WikiPathways WP560 TGF-β receptor signaling); (vi) myofibroblast/contractile differentiation (ACTA2, TAGLN, MYH11, MYL9, CNN1, POSTN; KEGG hsa04270 Vascular smooth muscle contraction, WikiPathways WP1991); and (vii) inflammatory-cytokine signaling (IL6, IL1B, CXCL8, CCL2, TNF, PTGS2; KEGG hsa04060 Cytokine–cytokine receptor interaction, WikiPathways WP530 Cytokines and inflammatory response). These gene sets were not exported verbatim from a single database; they were hand-curated from the established literature on fibroblast wound healing, with the individual gene membership mapped to the KEGG ^29, 30^ and WikiPathways ^31^ pathways listed above and to the corresponding Gene Ontology biological-process terms for wound healing, angiogenesis and ECM organization. The wound-healing-specific human maps WikiPathways WP5055 Burn wound healing, together with the bone-formation maps WikiPathways WP474 Endochondral ossification and WP4787 Osteoblast differentiation and related diseases relevant to bone healing, span several of these arms.

Note that these are culture-expanded PDL cells, not healing wounds. The analysis measures the intrinsic expression of wound-healing-associated gene programs, which is a molecular proxy for regenerative capacity — not a direct measurement of clinical healing speed.

The design is paired: each of the 4 donors (Sp1, Sp3, Sp5, Sp12) contributed one maxilla and one mandible sample. All statistics use the donor as the unit of replication (n=4), which is the correct, conservative approach and avoids treating 43,423 individual cells as independent replicates. With only 4 donors, statistical power is limited; absence of significance does not prove absence of a biological effect, but it does mean this dataset provides no positive support for the anecdote.

### These data do NOT support faster healing capacity in the maxilla

No wound-healing module differed significantly between jaws (all adjusted p > 0.75). Donor-to-donor variation dominated every module, and the direction of the (non-significant) jaw effect was inconsistent across donors — the paired donor lines cross repeatedly (Figure 9). If anything, the ECM/collagen/myofibroblast modules trended slightly HIGHER in the mandible, the opposite of the anecdote, while proliferation and angiogenesis trended marginally higher in the maxilla. None of these trends were statistically significant.

**Figure 9.**
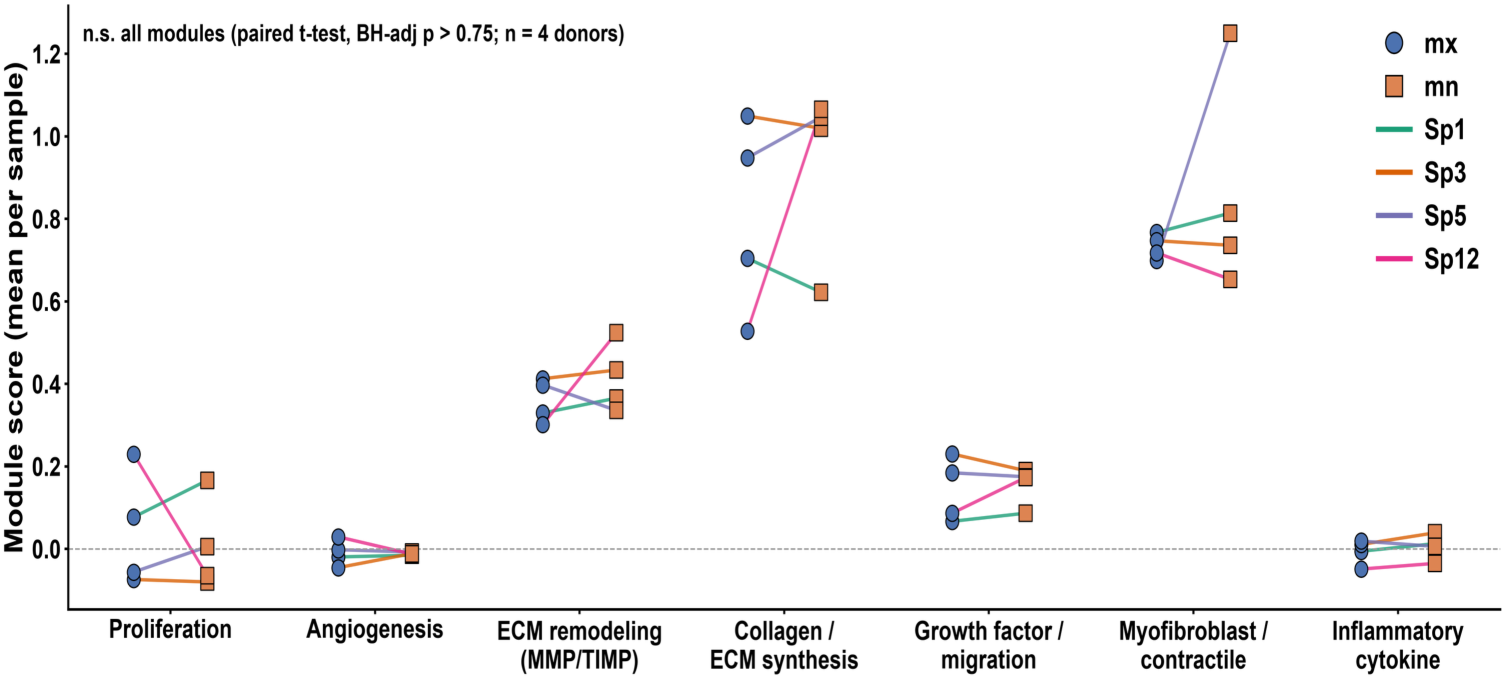
Per-donor paired wound-healing module scores, maxilla versus mandible. For each of seven wound-healing gene modules (x-axis), the mean per-sample module score is plotted for both jaw samples of every donor. Blue circles = mx (maxilla); orange squares = mn (mandible). Colored lines connect the paired mx→mn samples of the same donor (Sp1 green, Sp3 orange, Sp5 purple, Sp12 magenta); the dashed grey line marks a score of 0. The connecting lines change direction from donor to donor, indicating no systematic jaw effect (paired t-test, Benjamini–Hochberg-adjusted p > 0.75, n = 4 donors). Abbreviations: ECM, extracellular matrix; MMP, matrix metalloproteinase; TIMP, tissue inhibitor of metalloproteinases; mx, maxilla; mn, mandible.

**Table 2.** Module-level comparison (mean score across 4 donors)

| Module | Mean (maxilla) mx | Mean (mandible) mn | Higher in | Adj. p |
| --- | --- | --- | --- | --- |
| ECM remodeling (MMP/TIMP) | 0.360 | 0.415 | mn/mandible | 0.75 (n.s.) |
| Collagen / ECM synthesis | 0.807 | 0.938 | mn/mandible | 0.75 (n.s.) |
| Myofibroblast / contractile | 0.733 | 0.863 | mn/mandible | 0.75 (n.s.) |
| Inflammatory cytokine | -0.006 | 0.006 | mn/mandible | 0.75 (n.s.) |
| Proliferation | 0.044 | 0.007 | mx/maxilla | 0.82 (n.s.) |
| Growth factor / migration | 0.142 | 0.156 | mn/mandible | 0.82 (n.s.) |
| Angiogenesis | -0.010 | -0.012 | mx/maxilla | 0.89 (n.s.) |

### Individual wound-healing genes (from paired DESeq2, ∼donor + jaw)

Five wound-healing-relevant genes reached statistical significance in the gene-level analysis. The single result consistent with the anecdote is KDR (VEGFR2, angiogenesis receptor), higher in the maxilla. The others point the other way. Interestingly, Toll-like receptor 4 (TLR4) was higher in the mandibular PDLs. This finding is consistent with intrinsically stronger responses of mandibular PDLs to bacterial infection, because TLR4 signaling triggers pro-inflammatory cytokine production initiated by lipopolysaccharide (LPS) recognition, which leads to immune cell activation and inflammation. Moreover, an in vitro model of simulated orthodontic tooth movement demonstrated that TLR4 mediates inflammatory cytokine production upon mechanical compression corroborating the role of TLR4 in regulating mandibular inflammation ^6^.

**Table 3.** Wound-healing-relevant genes.

| Gene | log2FC (mn/mx) | Adj. p | Higher in |
| --- | --- | --- | --- |
| KDR | -1.95 | 0.048 | mx (maxilla) — angiogenesis |
| TLR4 | 3.91 | 0.000 | mn (mandible) — innate immune |
| GFRA1 | 1.35 | 0.000 | mn (mandible) |
| ACE | 1.20 | 0.028 | mn (mandible) |
| ERBB4 | 1.14 | 0.041 | mn (mandible) |

### Positional origin does not regulate MSC/PDL cell-program identity

Because craniofacial positional-identity transcription factors were the dominant jaw-associated signal, we asked whether positional origin also regulates the mesenchymal and periodontal-ligament (PDL) differentiation programs used to characterize these cells (Figure 8). Each of the six MSC/PDL programs (ISCT MSC-positive and MSC-negative markers, PDL/dental-MSC, osteogenic, adipogenic and chondrogenic) and the composite MSC/PDL identity index [(ISCT MSC⁺ + PDL/dental-MSC)/2 − ISCT MSC⁻] was tested for association with jaw of origin. Because the design is paired, the donor is the experimental unit; we therefore used per-donor paired t-tests (n = 4 pairs) as the primary test, with a cell-level linear mixed model (jaw as fixed effect, donor as random intercept; ∼43,000 cells) as a higher-power supplement. No program, and neither the composite identity, differed significantly between maxilla and mandible in the donor-paired analysis (all Benjamini–Hochberg-adjusted p > 0.33; composite identity Δ = +0.004, p = 0.94; Figure 10). The nominally largest signal — a lower adipogenic score in mandible — did not survive correction (raw p = 0.048, adjusted p = 0.33) and was not supported by the paired signed-rank test (p = 0.125). The per-donor trajectories for the composite score crossed rather than shifting in a common direction (Figure 10A), indicating that the sign of any jaw effect is donor-dependent rather than origin-dependent. Although the cell-level mixed model returned significant p-values for every program, this reflects the ∼43,000-cell count — pseudo replication of non-independent cells within each sample — rather than reproducibility across donors: all standardized effect sizes were small (|Cohen’s d| < 0.30; composite d = −0.06). Positional origin therefore does not detectably regulate the intrinsic MSC/PDL program identity of culture-expanded PDL cells in this cohort, although with only four donors the paired comparison is underpowered to exclude small effects.

**Figure 10.**
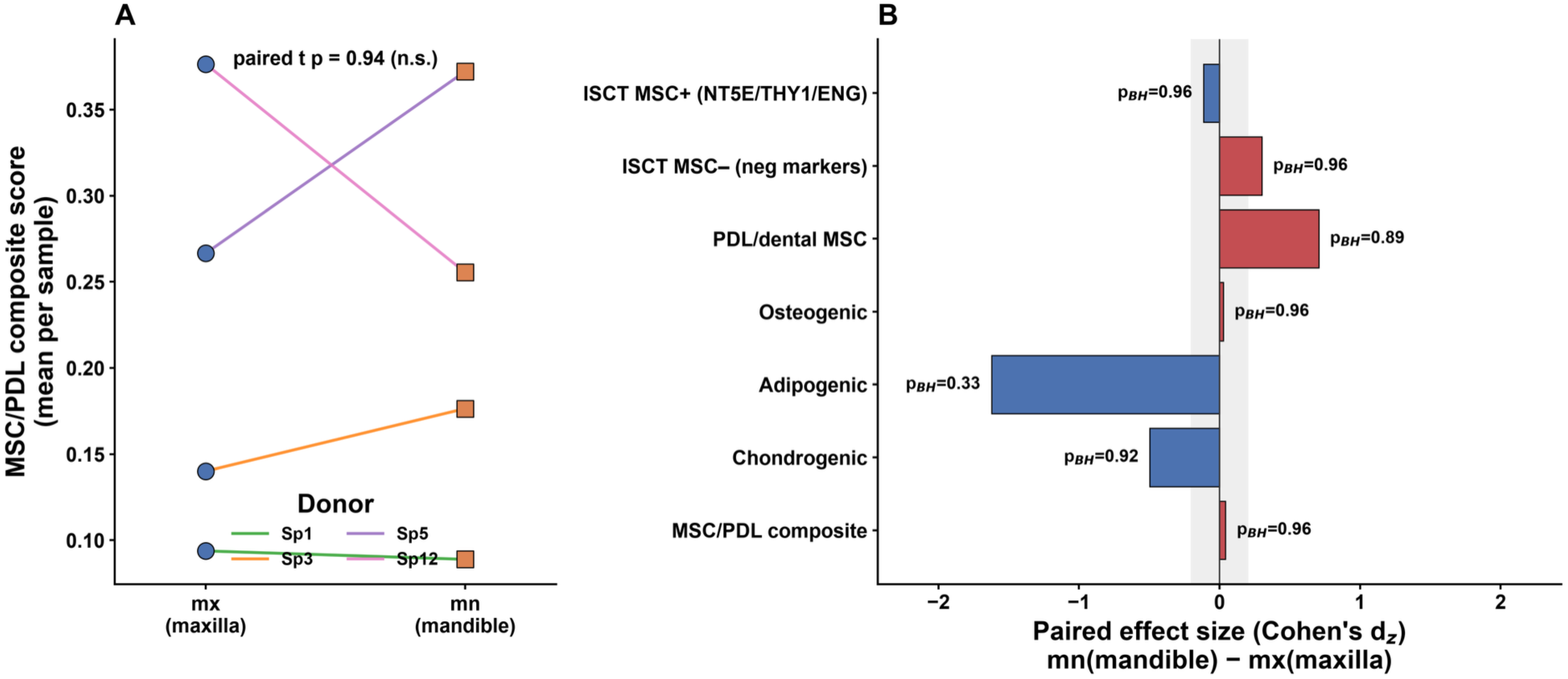
Positional origin (maxilla vs mandible) is not associated with MSC/PDL cell-program scores or composite identity. (A) Per-donor mean MSC/PDL composite-identity score for the maxilla (mx) and mandible (mn) sample of each donor; lines connect the paired samples of one donor (color) and cross rather than shifting consistently. The composite index is (ISCT MSC⁺ + PDL/dental-MSC)/2 − ISCT MSC⁻. Paired t-test p = 0.94. (B) Paired effect size (Cohen’s dᴢ, mandible − maxilla) for each of the six MSC/PDL programs and the composite identity; bars are colored by direction and the grey band marks the trivial-effect region (|dᴢ| < 0.2). Benjamini–Hochberg-adjusted paired-t p-values are annotated; none reached significance (all adjusted p ≥ 0.33; n = 4 donors). mx, maxilla; mn, mandible; ISCT, International Society for Cellular Therapy; dᴢ, paired Cohen’s d.

## Discussion

The single-cell transcriptomic survey presented here provides, to our knowledge, the first paired maxillary–mandibular scRNA-seq characterization of expanded human PDL cells across multiple donors. The resolution into distinct cell clusters illustrates donor variation and technical issues despite identical handling and expansion of cell isolates. This well-known phenomenon detracts from any bulk analysis yet can be resolved by single cell analysis. Three principal observations emerge. First, despite substantial numbers of profiled cells (43,423) and ten transcriptionally resolvable clusters, the expanded population was uniformly mesenchymal, with no detectable endothelial or immune lineage contribution. Second, the ten clusters corresponded not to discrete cell types but to functional fibroblast-like substates spanning proliferative, contractile, ECM/collagen-synthesizing, and progenitor-like transcriptional programs, several of which were strongly donor-restricted. Third, pseudobulk comparison of maxilla versus mandible cells, after accounting for donor as a blocking factor, revealed a modest but striking signature dominated by craniofacial positional-identity transcription factors (PITX1, BARX1, ALX1, NKX6-1), while wound-healing gene modules showed no significant jaw-associated difference. Taken together, these findings position cultured PDL cells as retaining developmental positional memory while largely losing the cellular and transcriptional complexity of the native tissue niche. Positional origin was found to determine pathological calcification of adult vascular smooth muscle cells ^32^. Here, we found no evidence that positional origin regulates PDL cell-program identity: across the six mesenchymal and periodontal-ligament differentiation programs and the composite MSC/PDL identity index, none differed significantly between maxilla and mandible in a donor-paired analysis (all adjusted p > 0.33; Figure 10), and the direction of any per-donor difference was inconsistent between individuals. Thus, unlike the pathological calcification of vascular smooth muscle cells, the intrinsic MSC/PDL differentiation programs of culture-expanded PDL cells appear not to carry a detectable maxillary–mandibular positional signature, even though these same cells retain positional-identity transcription-factor expression — though with four donors we cannot exclude small effects.

The absence of endothelial and immune signatures in every cluster stands in clear contrast to scRNA-seq studies of native or minimally manipulated periodontal tissue, which consistently resolve PDL, gingiva, and epithelium into multiple lineages, including clusters bearing immune-response gene signatures alongside mesenchymal populations ^21^. Similarly, tissue-level transcriptomic profiling of PDL has shown that collagens, proteoglycans, and structural ECM genes characteristic of the native matrix niche are markedly attenuated once cells are placed into monolayer culture, with canonical PDL markers such as osteopontin, asporin, periostin, and osteonectin all downregulated relative to intact tissue ^3^. Similarly PLAP-1/Asporin, which is characteristically abundant in native PDL tissue ^28^) was detected in only 4.3% of all cells, consistent with the known tendency of ex-vivo expansion to attenuate tissue-specific matrix programs. Our data extend this observation to the single-cell level: not only are matrix and marker genes reduced on average, but the entire non-fibroblastic cellular ecosystem — vascular, perivascular, and immune populations that are integral to PDL homeostasis and remodeling in vivo — is absent by the time cells have been expanded for scRNA-seq. This is consistent with the general expectation that ex vivo expansion selects for a mesenchymal, proliferation-competent population while diluting or eliminating minority lineages that lack robust growth capacity under standard culture conditions, reinforcing the view that cultured fibroblast preparations represent an artificial, immature cellular state rather than a faithful surrogate for the tissue of origin ^3^. This contrast is made explicit by native PDL single-cell atlases, in which lineage tracing and transcriptomic profiling of intact murine PDL recover not only fibroblastic PDL cells but also Nes+ mural cells, S100B+ Schwann cells, and CD45+, CD31+, and Epcam+ immune, endothelial, and epithelial populations—precisely the non-mesenchymal compartments that are absent from our expanded cultures ^23^. PDL organoid culture may help to restore and maintain organotypic gene expression. Nonetheless, within this uniformly mesenchymal population, the substates we identified — proliferating, collagen-high, ECM/decorin, contractile (MYLK/PDE5A), VCAN+ progenitor-like, stress/MT-high, and donor-restricted clusters — align conceptually with prior reports of PDL cellular heterogeneity, even though the specific cluster identities differ across platforms and preparations. Native-tissue scRNA-seq has similarly resolved PDL mesenchymal populations along matrix-composition axes, with Mkx- and Scx-expressing subsets distinguished by proteoglycan/elastic-fiber versus collagen I/III programs and has shown that loss of the Mkx regulator shifts the balance toward ossification-prone and inflammatory phenotypes ^19^. Likewise, functionally distinct PDL fibroblast subsets producing IL-1β versus RANKL have been described in inflamed periodontal tissue, underscoring that fibroblast heterogeneity is not merely descriptive but carries distinct pathophysiological consequences ^20^. Clone-based studies provide an orthogonal line of evidence for this heterogeneity at the progenitor level: single-cell-derived PDL clones segregate into osteoblastic/cementoblastic versus fibroblastic phenotypes with distinguishing Wnt/cadherin signatures ^10^, and Zbp1 expression marks a subset of clones with disproportionately high osteogenic capacity despite indistinguishable morphology and standard marker expression ^11^, although the corresponding transcript was essentially undetectable in our culture-expanded human PDL cells (SFigure 2). Our VCAN+ progenitor-like and contractile cluster 6 (Figure 7, 8) may represent culture-compatible cell populations of these previously defined osteogenic- or myofibroblastic-primed substates, consistent also with single-cell reports that hPDLSC cultures resolve into osteogenic and myofibroblastic-dominant populations ^17^ and that cultured PDLSCs show lower proliferative fractions and distinct trajectory branching relative to dental pulp stem cells ^18^. Cluster 6 VCAN+ progenitor-like and contractile cells thus come closest to putative PDLSC. They represent 15.4% total cells. VCAN/versican-based cell selection and complex culture technique combining PDL cells with stem cell niche cells into organoids may help to derive sufficient PDLSC for therapeutic applications.

Because these clone- and tissue-level studies implicate specific transcription factors in PDL identity and lineage priming, we examined the expression of the PDL/tendon-associated transcription factors MKX (mohawk), SCX (scleraxis) and ZBP1 directly in our culture-expanded cells (SFigure 2). MKX was broadly retained, detected in 61.4% of cells and expressed most strongly along the extracellular-matrix axis — highest in the ECM-high, ECM/decorin and collagen-high fibroblast states and reduced in the proliferating fractions — mirroring the Mkx-associated proteoglycan and collagen matrix programme described in native PDL ^19^. In contrast, SCX was largely absent (1.2% of cells, at most two transcripts per cell) and ZBP1, which marks high-osteogenic murine PDL clones ^11^, was essentially undetectable (8 transcripts across all 43,423 cells). Thus, of the PDL/tendon-associated transcription factors examined, only MKX survived monolayer expansion at appreciable levels while retaining state-specific structure, whereas SCX and ZBP1 were strongly attenuated (SFigure 2) — consistent with the broader downregulation of tissue-specific programmes upon ex-vivo culture and reinforcing the view that expanded PDL fibroblasts represent an immature, partially de-differentiated state relative to their tissue of origin.

A complementary view of PDL identity comes from the cementoblast-versus-ligament-fibroblast axis, along which human PDL cells can be directed into transcriptionally distinct membrane and extracellular-matrix programmes ^12^. Examining these markers in our cultures (SFigure 1), the cementoblast marker CEMP1 (cementum protein 1) was essentially undetectable (0.08% of cells, 35 transcripts across all 43,423 cells), whereas PLAP-1/asporin — a characteristic PDL/ligament matrix proteoglycan — was detected in 4.3% of cells and significantly enriched in the ECM/decorin and proliferating fibroblast states. Unfortunately, given the design of this scRNAseq study, cementum attachment protein (CAP) is not an independent gene but a splice variant of HACD1/PTPLA (3-hydroxyacyl-CoA dehydratase 1/protein tyrosine phosphatase-like): the CAP transcript (GenBank AY455942.1) encodes a truncated 140-amino-acid polypeptide (AAR22554.1) whose N-terminal 125 residues match a short PTPLA isoform but carry a proline in place of the catalytic arginine, and this PTPLa/CAP form is expressed in cementoblasts and probable cementoblast precursors (Montoya et al., 2014). Our scRNA-seq reference however, annotates HACD1 as a single gene-level feature and does not resolve this splice variant, so HACD1 counts — broadly detected in 44.3% of cells across all states, consistent with its housekeeping role in fatty-acid elongation — pool authentic HACD1 with any CAP-variant reads and cannot be equated with CAP; a CAP-specific measurement would require isoform-level resolution of the retained-intron junction that defines the variant, which these gene-level data do not provide. Until true CAP transcript expression is confirmed we posit that the retention of a ligament-type matrix marker together with the near-absence of a cementoblast marker indicates that base-medium expansion yields cells resembling the ligament-fibroblastic rather than the cementoblastic progenitor state, consistent with the divergent differentiation trajectories and matrix repertoires reported for these two lineages ^12^.

That contractile and mechanoresponsive transcriptional programs persist in our expanded cells is also consistent with the long-established role of PDL fibroblasts as mechanosensitive cells ^33^ that couple integrin-matrix engagement to cytoskeletal remodeling and force-dependent tissue maintenance in vivo ^1^, a property that is functionally reinforced by in vitro work showing that mechanical compression and growth-factor stimulation (FGF1) modulate proliferation, osteogenic marker induction, and inflammatory cytokine output in cultured PDL fibroblasts ^15^. Consistent with our resolution of multiple fibroblast substates within an in vitro-expanded population, single-cell analysis of cultured PDL- and gingiva-derived MSCs has similarly identified both tissue-specific and tissue-spanning subpopulations bearing distinct upregulated gene sets, confirming that culture-expanded PDL preparations retain intercellular transcriptional heterogeneity ^22^.

The jaw comparison offers a more nuanced picture than might have been anticipated from prior functional studies. We found that the dominant, statistically robust transcriptional distinction between maxillary and mandibular PDL cells was not a difference in proliferative or osteogenic gene programs but the retained expression of positional-identity transcription factors — PITX1, higher in the mandible, has a well-established role in lower-jaw (first pharyngeal arch) identity, where its mesenchymal expression is required for normal mandibular development ^34, 35, 36^. The maxilla-enriched factors we detected — BARX1, ALX1 and NKX6-1 — are craniofacial or developmental patterning genes, but their upper-jaw directionality here is an observation in adult culture-expanded PDL cells rather than a previously established maxillary positional code: BARX1 is expressed in the mesenchyme of both arches and is more closely associated with molar/proximal identity, and the ALX family marks frontonasal and upper-midface mesenchyme, so the jaw-side assignment of these candidate factors awaits developmental confirmation. This suggests that expanded PDL cells retain a form of epigenetic or transcriptional “positional memory” of their jaw of origin even after prolonged in vitro passage, a phenomenon broadly analogous to the persistence of HOX-code identity in fibroblasts ^37^, which has been termed the “ZIP-code” identifying original body sites ^38^. The retention of craniofacial patterning transcription factors is developmentally plausible given that the PDL arises from cranial neural crest-derived dental follicle mesenchyme, whose lineage diversification is governed by spatially organized, cell-type-specific gene regulatory networks during tooth morphogenesis ^39^; the positional-identity signature we observe may therefore represent the persistence of a developmentally established, lineage-intrinsic transcriptional program rather than an artifact of culture.

Notably, the PDL positional signature was not accompanied by a broader divergence in proliferation- or differentiation-associated transcriptional programs, which contrasts with functional studies reporting that maxillary PDLSCs proliferate faster and differentiate more robustly along osteogenic and chondrogenic lineages than mandibular PDLSCs ^7^, and with kinomic profiling showing intrinsic differences in kinase activity — including stronger EphA signaling in mandibular cells and greater PI3K-Akt pathway activity favoring mandibular osteogenic modulation — between paired maxillary and mandibular PDLSCs from the same donors ^24^. One interpretation is that the site-specific functional differences reported by Mert and Malyaran operate largely at the level of protein activity, kinase signaling, and post-transcriptional regulation rather than steady-state transcript abundance, meaning that our transcriptomic assay — while sensitive enough to detect subtle, biologically meaningful patterning-gene differences — may simply not be well positioned to capture the signaling-level heterogeneity that drives functional divergence in proliferation and differentiation assays.

Despite the persistent clinical impression that maxillary periodontal tissue heals faster than mandibular tissue, we found no significant difference in wound-healing-associated gene modules between jaws (all adjusted p > 0.75), with the qualified exception of higher KDR/VEGFR2 expression in maxillary cells, consistent with a modestly greater angiogenic priming. We interpret this negative result cautiously rather than as evidence against a true biological jaw difference in healing capacity. Clinical wound healing is an emergent, multicellular, and multi-tissue process involving vasculature, immune cell recruitment, and dynamic matrix remodeling under mechanical and inflammatory cues that are entirely absent from an expanded, monoculture, immune- and endothelial-cell-free in vitro system. Functional assays that directly interrogate osteogenic differentiation, mineralization, and osteoclast-inductive capacity — the kind of assays that revealed superior maxillary proliferative and osteogenic potential ^7^ and jaw-specific kinase signaling ^24^ — may be inherently more sensitive to site-specific regenerative differences than a steady-state transcriptomic snapshot of cultured cells, in the same way that PDL cells only reveal their comparatively greater osteogenic and osteoclast-inductive activity relative to adjacent alveolar bone cells when assayed functionally rather than by static transcriptional profiling ^14^. It is also plausible that healing-relevant differences are conferred non-cell-autonomously by local vascular density, innervation, or immune microenvironment rather than by intrinsic PDL fibroblast transcriptional programs, in which case no amount of refinement of a fibroblast-only assay would be expected to recover the clinical phenotype. In conclusion, in this PDL single-cell dataset, wound-healing-associated transcriptional programs were indistinguishable between maxilla and mandible cells once donor identity was accounted for. The data neither supported nor strongly refuted the clinical anecdote of faster maxillary healing; they simply showed no intrinsic, cell-autonomous transcriptional advantage in either jaw at the level measured. A faster clinical healing in the maxilla, if real, would more likely be driven by anatomical and physiological factors (vascular supply, bone density/type, mechanical load, saliva exposure) than by an intrinsic difference in the PDL fibroblast transcriptome.

Several limitations temper the interpretation of these findings. Most importantly, the cells analyzed were expanded in monolayer culture prior to sequencing, and culture expansion is well documented to suppress key PDL marker genes and to shift cells toward a more proliferative, immature transcriptional state relative to native tissue ^3^; our uniformly mesenchymal, fibroblast-substate composition should therefore be understood as a property of the cultured population rather than a claim about native PDL tissue composition, which is known to be considerably more heterogeneous, including immune-associated clusters, at the single-cell level ^21^. The donor cohort was small (n = 4), and donor identity — rather than jaw of origin — was the dominant source of transcriptional variance, with several clusters being almost entirely restricted to individual donors (e.g., cluster 2 at ∼86% Sp1; clusters 3 and 7 largely Sp5/mandible-restricted); this donor-dominated structure limits the generalizability of both the substate catalog and the jaw-comparison findings and raises the possibility that some “jaw” differences partly reflect unbalanced donor representation rather than pure anatomical signal. Cluster annotation was based on marker-gene inference and known functional gene sets rather than functional validation (e.g., lineage tracing, clonal assays, or differentiation challenge), so the proliferating, contractile, ECM-high, and progenitor-like labels should be regarded as descriptive hypotheses rather than confirmed cell states, in contrast to the functionally validated osteogenic/fibroblastic clone distinctions established by direct differentiation assays ^10, 11^. Finally, our pseudobulk DESeq2 design, while appropriately conservative in modeling donor as a blocking factor, is likely underpowered to detect all but the largest and most consistent jaw-associated effects, which may explain why only a small, patterning-gene-dominated set of transcripts reached significance. This donor-dominated structure contrasts with a single-cell study of cultured PDL- and gingiva-derived MSCs that reported donor-to-donor transcriptional differences to be statistically significant but limited in magnitude and confined to specific cellular processes ^22^; the discrepancy may reflect our smaller and more unbalanced donor cohort, differences in culture and dissociation protocols, or the per-donor clustering strategy employed in that study, and it underscores the need for larger, balanced cohorts before the relative contributions of donor and anatomical site can be resolved.

These findings carry both cautionary and constructive implications for PDL regenerative research. They reinforce that transcriptomic profiling of expanded cell populations — however granular at single-cell resolution — cannot substitute for functional and in vivo assays when the question at hand is regenerative or healing capacity, echoing evidence that inflammatory and signaling-level modulation of PDLSC differentiation, such as CB1-receptor-mediated rescue of TNF-α/IFN-γ-impaired osteo/dentinogenic differentiation via p38 MAPK and JNK signaling ^16^, is not necessarily reflected in steady-state transcript abundance yet is clearly biologically consequential. At the same time, the retained expression of craniofacial positional-identity transcription factors after extended culture is a notable and, to our knowledge, previously unreported finding that merits mechanistic follow-up, including chromatin-level studies to determine whether this positional memory is maintained through persistent enhancer activity or epigenetic marking established during development. Future work should prioritize paired functional assays (proliferation, osteogenic/chondrogenic differentiation, osteoclast induction) alongside transcriptomics in the same donor-matched maxillary/mandibular samples, larger and more balanced donor cohorts to disentangle donor from anatomical effects, and, where feasible, direct single-cell profiling of freshly isolated or minimally cultured PDL tissue to determine which of the substates and jaw-associated signatures described here are intrinsic to PDL cells versus artifacts of ex vivo expansion. Such integrative approaches will be necessary to reconcile the modest transcriptional jaw differences observed here with the more pronounced functional and clinical asymmetries reported in the literature.

## Acknowledgments

This work was funded by the Deutsche Forschungsgemeinschaft (DFG, German Research Foundation – TRR219 – Project-ID 322900939, (NM, WJ-D), SFB 1739 - Project-ID 546544928 (MW, NM, RBC, IK), DFG-Project-ID 403041552 (WJ-D), 490932300 (SN-S, MW, RBC), 504777725, 559483338, 583318578 (MW, RBC). This work was supported by the Genomics Facility of the Interdisciplinary Center for Clinical Research (IZKF) within the Faculty of Medicine at RWTH Aachen University. Publication charges were covered by project DEAL.

## Notes

### Competing Interest Statement

The authors have declared no competing interest.

